# Feasability of Baseline cardIac investiGation in awake elepHants (*Elephas mAximus*) using tRansThoracic echocardiography: the BIG HEART study

**DOI:** 10.1101/2025.09.16.676569

**Authors:** Antonin Boutibou, Valérie Chetboul, Mathieu Magnin, Morgan Bureau, Coline Baillon, Norin Chai

**Author notes:** Corresponding author: Prof. Valérie Chetboul.

## Abstract

**Introduction:** Cardiovascular diseases are relatively common in elephants, but most cases are diagnosed only postmortem. Transthoracic echocardiography (TTE) is widely used in domestic and captive animals, although its application in elephants has not yet been established. The aim of this pilot study was to determine the feasibility of performing TTE in Asian elephants (*Elephas maximus*), to develop a standardized procedure to optimize image quality, and to assess measurement variability.

**Materials and Methods:** Preliminary trials were conducted in ten elephants, including seven free-ranging animals in Cambodia and three captive elephants in France. These led to a standardized TTE protocol based on cooperative training and positive reinforcement. Key refinements included the exclusive use of a left parasternal approach, soaking the thoracic skin with lukewarm water before application of coupling gel, and extension of the left forelimb over a training ball to optimize access to the acoustic window. Each examination included two LV views: a long-axis view for M-mode measurements and a short-axis view for two-dimensional (2D) analysis. A total of 72 examinations were performed on 4 days on three elephants, with offline assessment of eight 2D and M-mode variables, including two indices of LV function (shortening fraction and fractional area change). A general linear model was used to determine within-day and between-day coefficients of variation.

**Results:** All examinations were successfully completed without anesthesia or restraint. Within-day and between-day variability were low for all parameters (coefficients of variation <10%), while interindividual variability was higher (5%-19.6%).

**Discussion:** LV dimensions and systolic function can be assessed in awake Asian elephants with excellent repeatability and reproducibility.

**Conclusions:** The validated TTE procedure developed here enables longitudinal cardiac monitoring in captive Asian elephants without anesthesia, providing a unique opportunity for the antemortem diagnosis of left-sided heart disease and facilitating future comparative and conservation studies in this endangered species.

## Introduction

Cardiovascular diseases are increasingly reported as an important cause of morbidity and mortality in elephants, affecting both Asian elephants (*Elephas maximus*), listed as Endangered, and African elephants (*Loxodonta africana* and *Loxodonta cyclotis*), listed as Vulnerable and Critically Endangered respectively, according to the IUCN Red List [1–3]. Various cardiovascular disorders have been identified in these species, some of which resulting in sudden death or the development of congestive heart failure. Documented noninfectious acquired conditions include dilated and hypertrophic cardiomyopathies, as well as arteriosclerotic changes primarily affecting the aorta, coronary arteries, and major aortic branches [1,2,4]. In addition, viral myocarditis cases associated with Picornaviridae or West Nile virus infections have also been described [1,4]. Several retrospective necropsy surveys have highlighted the importance of cardiovascular pathology in both Asian and African elephants [1,2]. In a survey of 379 captive elephants, cardiovascular disease was documented in only 5% of cases, but 95% of these animals died, with cardiac pathology directly responsible for 61% of the deaths [1]. In a more recent study analyzing 336 necropsy reports from the European Association of Zoos and Aquaria, cardiovascular diseases represented the leading category of lesions in African elephants (24% of cases) and the third most common category in Asian elephants (15% of cases), after infectious diseases (notably, elephant endotheliotropic herpesvirus, EEHV) and musculoskeletal disorders [2]. Such data underline both the clinical importance of cardiac disorders in elephants and the fact that most knowledge to date has been derived from necropsy rather than functional investigations in living animals.

In human and veterinary cardiology, in both domestic and captive animals, transthoracic echocardiography (TTE) is the gold standard non-invasive imaging modality for assessing cardiac anatomy and function, allowing real-time evaluation of chamber dimensions, wall thicknesses, myocardial contractility, and valve assessment, thereby facilitating early recognition of pathological alterations and guiding therapeutic interventions [5]. Despite its widespread application across numerous animal species, echocardiographic data remain extremely limited in elephants, and its clinical use is still rare. The paucity of such data can be attributed not only to the large body size but also to the particularly thick skin and subcutaneous tissues, and to the cranial and relatively deep position of the heart within the thorax, which limit acoustic access [1]. Additional logistical and ethical constraints also contribute to this scarcity, as echocardiographic evaluation requires close physical proximity to the thorax of these massive animals, ideally in standing, awake conditions to avoid the confounding effects and risks of anesthesia, but with inherent safety concerns for operators. To the best of our knowledge, only fragmented echocardiographic observations have been reported to date, with no standardized protocol or population-based reference values available to guide interpretation.

Given the prevalence and impact of cardiac diseases in elephants, establishing a repeatable, reproducible, and non-stressful TTE procedure is essential for preventive and diagnostic strategies. Additionally, infection with EEHV, a major cause of mortality in young Asian elephants, targets endothelial cells and can induce severe vascular dysfunction and myocardial injury [6,7]. Reliable antemortem cardiac assessment tools, such as TTE, are therefore needed to detect such complications that are most often recognized only postmortem.

The aims of the present pilot study were therefore 1) to determine the feasibility of performing TTE in awake, apparently healthy Asian elephants (*Elephas maximus*), 2) to develop a standardized and non-stressful TTE procedure to optimize image quality, and 3) to assess the intra-observer within-day (repeatability) and between-day (reproducibility) variability of the corresponding measurements.

## Materials and Methods

### Animals

Faisability echocardiographic trials were conducted in ten Asian elephants (*Elephas maximus*), including seven free-ranging animals in Cambodia in collaboration with the Elephant Valley Project (the same cohort used by our group to develop an electrocardiogram (ECG) recording method in this species [8]), and three captive elephants from La Tanière - Zoo Refuge (Nogent-le-Phaye, France). The latter three elephants were included in the validation protocol of the present study. They were males, apparently healthy as determined by routine veterinary monitoring, aged 7, 9, and 13 years, with shoulder heights of 2.02, 2.42, and 2.51 m, and body weights of 1,800, 2,600, and 3,600 kg, respectively (EL1, EL2, and EL3).

The facilities housing the three study elephants included one indoor enclosure, subdivided into four small connected areas, with controlled access to two outdoor areas. The indoor and outdoor enclosures contained furnishings that added physical complexity to their environment (e.g., rocks, scratching poles, suspended food access). A staff of four keepers cared for the elephants, with at least two present on site each day. In addition, one full-time veterinarian was responsible for their health care and welfare. The elephants were fed twice daily (at 10:00 a.m. and 4:00 p.m.) with hay, forage, and browse. They also received daily enrichments (feeding, physical, and cognitive), using apples, bananas, mangos, and training biscuits (DK training biscuits, DK Zoological, Putten, Netherlands). Finally, the elephants participated in daily medical and husbandry training sessions based on operant conditioning techniques. When an elephant successfully performed the requested behavior, the keeper produced a whistling sound as a secondary reinforcer, followed by a reward (e.g., a piece of fruit) and verbal praise.

### Ethical considerations

The cardiac ultrasound examinations performed in this study were carried out without the use of anesthesia, as part of routine clinical care, on captive elephants regularly trained to voluntarily participate in veterinary procedures through positive reinforcement techniques. All elephants remained in their familiar environment throughout the whole study period. These animals undergo daily training sessions that enable them to accept medical procedures such as foot presentations for pedicures, oral inspections, and even blood sampling for preventive health assessments without the need for restraint or stress. Consequently, the echocardiographic examinations were fully integrated into an established health-monitoring program and did not involve any experimental design or intervention beyond standard clinical practice. They therefore cannot be considered animal experimentation or any additional intervention that could compromise animal welfare.

This type of cooperative training is now widely recognized as essential to the management of captive elephant health, serving both animal welfare and handler safety [9,10]. It avoids the use of invasive procedures, physical restraint, or anesthesia, while promoting voluntary cooperation during clinical interventions [11,12]. Moreover, such training is considered a form of cognitive and social enrichment, contributing positively to the animals’ emotional and physical well-being. Within this framework, the procedures described here fall strictly under the scope of routine veterinary practice, and thus did not require prior approval from an ethics committee according to experimental animal use regulations. They are also in line with international standards for the management of captive elephants, as emphasized by the Association of Zoos and Aquariums [12], which highlights voluntary training to minimize the need for anesthesia, and by the European Association of Zoos and Aquaria [13], which explicitly recommends cooperative training for diagnostic procedures such as ultrasound examinations.

### Transthoracic echocardiographic technique and measurements Development and feasibility of the transthoracic echocardiographic procedure (**Figs. 1 and 2**)

All TTE examinations were performed using a portable cardiovascular ultrasound system (Vivid iq, GE Healthcare, 9900 Innovation Drive, Wauwatosa, WI 53226, USA) equipped with a 4C-RS convex transducer (1.8-6 MHz). Our preliminary trials demonstrated that the 4C-RS probe facilitated identification of the heart - a challenging step given the size of the animal - and provided a wider field of view, particularly in the cardiac long-axis plane, compared with the 3S phased-array transducer (1.5-3.5 MHz). The 4C-RS transducer was therefore selected for this study (**Fig. 1A**).

**Fig 1.**
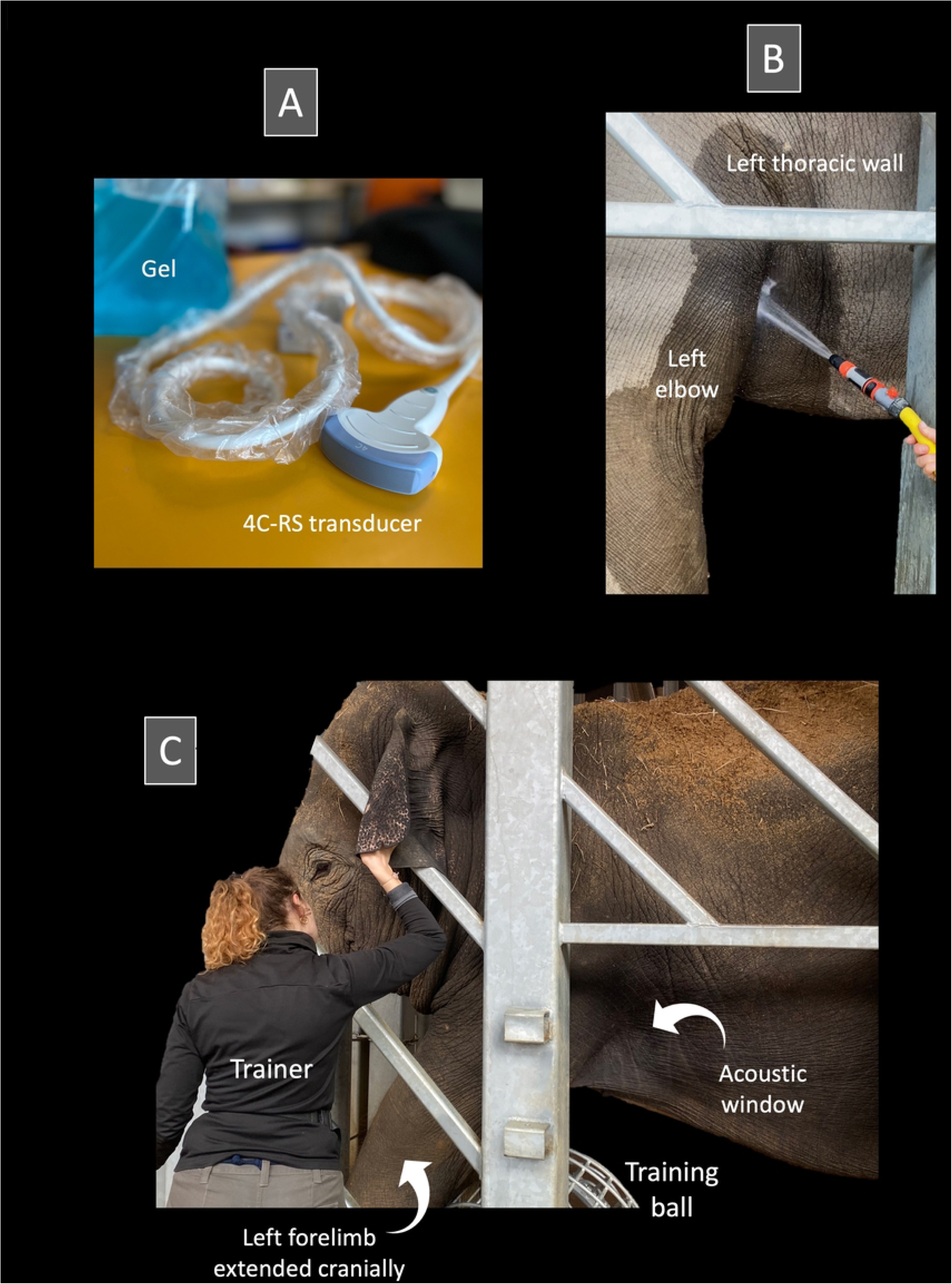
(A to C): Echocardiographic procedure in the elephants included in the study. **(A)** A 4C-RS convex transducer was used, which provided a wide ultrasound field suitable for cardiac imaging in this large species. **(B)** Prior to imaging, the acoustic window located slightly dorsal and caudal to the left elbow was rinsed with lukewarm water to soften the thick skin and improve acoustic coupling with the large amount of ultrasound gel applied, a step that was well tolerated and appeared to be positively perceived by the elephants. **(C)** The elephant stood calmly and without restraint against the protective barrier while the trainer gently interacted with its head. The left forelimb was extended cranially, facilitated by resting it over a training ball placed on the ground, in order to optimize access to the cranial portion of the left cranial thoracic wall and echocardiographic window. *Photos by Valérie Chetboul*.

**Figure 2.**
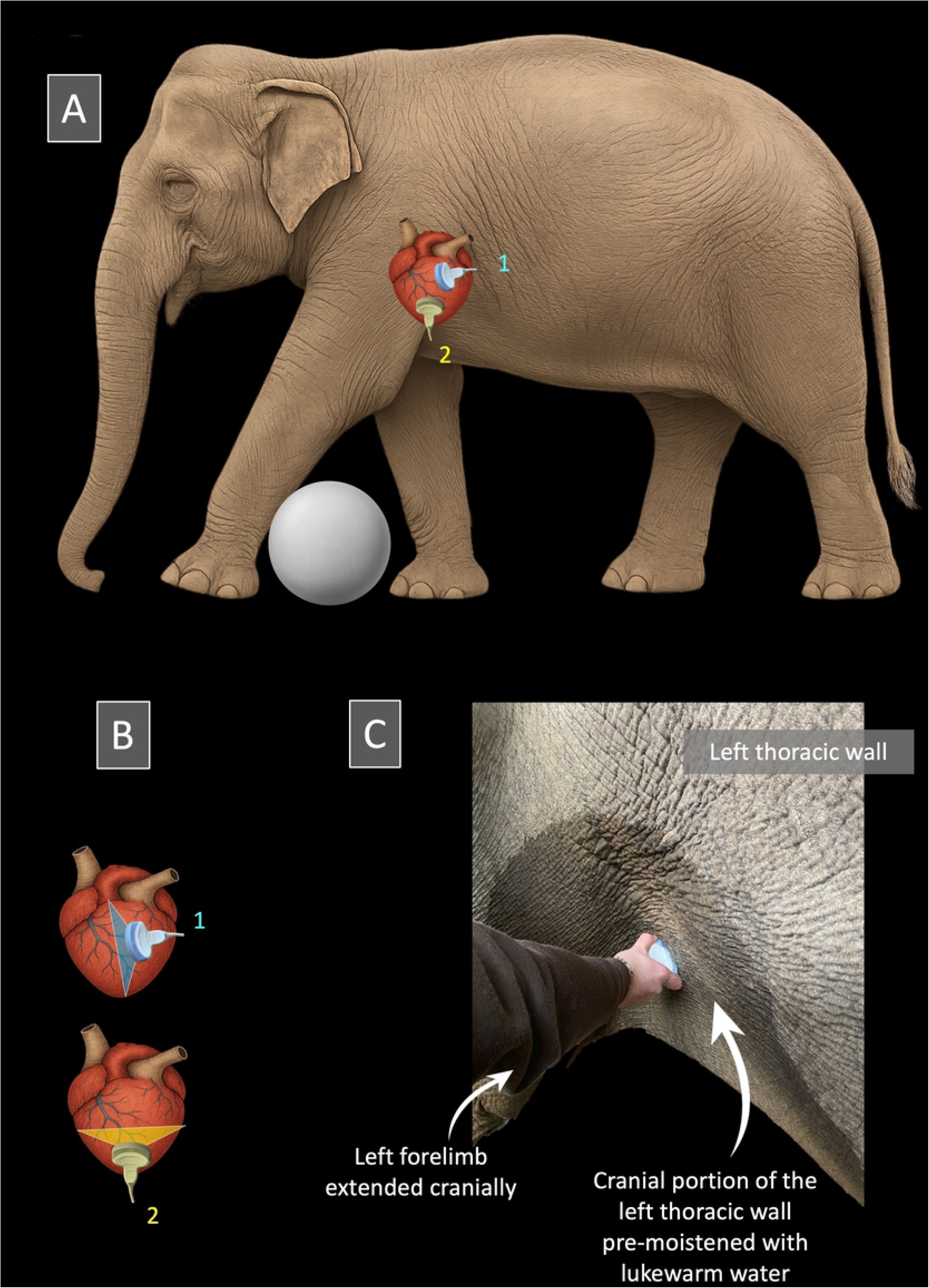
(A to C). Transducer positioning for echocardiographic imaging of the elephants included in the study. **(A) and (B)** Elephant positioning for transthoracic echocardiographic examination and probe placement to obtain the left parasternal long-axis view (1) and the left parasternal short-axis view at the level of the left ventricle (2). **(C)** Position of the transducer to obtain the left parasternal long-axis view. The probe is placed in an intercostal space located caudal and slightly dorsal to the left elbow, aligned parallel to the ribs. In both Figures 2A and 2C, the left forelimb is gently extended cranially, facilitated by resting over a training ball placed on the ground, in order to improve access to the cranial portion of the left cranial thoracic wall. For optimal imaging, this acoustic window was first moistened with lukewarm water to soften the skin, before application of a large amount of ultrasound gel, thereby enhancing ultrasound transmission. *Illustration and photo by Valérie Chetboul*.

Animal training for the TTE procedure was based on operant conditioning with exclusively positive reinforcement. The refinement of the technique initially consisted of identifying the optimal acoustic window through preliminary echocardiographic trials first conducted in seven free-ranging Asian elephants in Cambodia [8], and then in the three elephants included in the present validation study. Attempts to obtain right parasternal views - which represent the standard imaging approach in domestic carnivores and horses [5] - proved unsuccessful, as the heart was either not visible or only poorly visualized depending on the individual. Consequently, only the left parasternal approach was retained, with the probe positioned on the left hemithorax, caudal and dorsal to the left elbow, in an intercostal space. Using this left parasternal approach, the left ventricle (LV) could be consistently visualized in two-dimensional (2D) mode in both its long-axis and short-axis planes, hereafter referred to as the left parasternal long-axis view and the left parasternal short-axis view, respectively. In contrast, the right heart cavities, the aorta, and the left atrium were only poorly visualized with this approach and were therefore not measured.

Preliminary trials also revealed that image quality was improved by a two-step preparation of the acoustic window: first, the thick skin was soaked with lukewarm water, ensuring sufficient time for adequate softening (i.e., at least 3–5 minutes under water, followed by an additional 5 minutes of waiting), with then the application of a generous amount of coupling gel (**Fig. 1B**).

### Final transthoracic echocardiographic procedure (**Figs. 1 and 2**)

The three elephants included in the validation study were trained to remain standing for several minutes against the enclosure. They were then gradually desensitized to the preparation of the acoustic window, which involved soaking the skin with lukewarm water and then applying generous amounts of ultrasound coupling gel to the thorax. Finally, they were accustomed to the placement of the probe on their thorax without reacting to the procedure. After three months of training, the three elephants were readily conditioned to voluntarily maintain the optimal position for several minutes, allowing acquisition of the TTE images.

The final TTE method was as follows: the trainers were responsible for bringing the elephants to an appropriate location, as close as possible to the protective barrier (**Fig. 1C**). They also trained the elephants to move their left forelimb forward (using a large training ball as a target for positive reinforcement), in order to optimize access to the echocardiographic window located cranially, immediately caudal and slightly dorsal to the left elbow (**Fig. 1C**). While the trainers distracted the elephants by gently talking to them and offering food (apples), optimal acoustic windows were promptly identified and obtained by one observer (Observer 1, AB), while a second observer (Observer 2, VC, Diplomate of the European College of Veterinary Internal Medicine - Cardiology) simultaneously adjusted the imaging settings. The overall gain was set to its maximum to optimize visualization of the LV myocardial walls. All measurements were subsequently performed offline by Observer 2. Each examination systematically included the two aforementioned 2D views, i.e., the left parasternal long-axis view and the left parasternal short-axis view. The left parasternal long-axis view was used to position the M-mode cursor and obtain the transventricular M-mode view, allowing measurement of left ventricular walls and cavity dimensions in both diastole and systole, whereas the left parasternal short-axis view was used to calculate an index of radial systolic LV function, namely the LV fractional area change (LV FAC%).

### Left parasternal LV long-axis view and corresponding M-mode measurements (Figs. 2 and 3)

The left parasternal long-axis view was obtained by placing the transducer in the intercostal space immediately caudal and slightly dorsal to the left elbow, with the ultrasound beam oriented parallel to the ribs and fine adjustments made according to image quality (**Fig. 3**). This view displayed the LV cavity, bordered proximally by the LV free wall (LVFW) and distally by the interventricular septum (IVS). In some animals and examinations, a portion of the right ventricle was also visible in the lower part of the image (**Fig. 3**). The M-mode cursor was positioned perpendicular to the LV myocardial walls, immediately below the insertion of the papillary muscles, which are particularly well developed in elephants, in order to avoid including them in the measurements. M-mode echocardiograms showing at least two consecutive cardiac cycles were obtained for measurement of LV diameters and myocardial wall thicknesses (**Fig. 3**). Left ventricular end-diastolic and end-systolic internal diameters (LVIDd and LVIDs, respectively), LVFW thicknesses at end-diastole (LVFWd) and end-systole (LVFWs), IVS thicknesses at end-diastole (IVSd) and end-systole (IVSs) were measured on this view using the leading edge-to-leading edge technique, and the LV shortening fraction (SF%) was then calculated. As no concomitant ECG tracing could be performed, end-diastole and end-systole were defined based on mechanical events; specifically, end-diastole was identified as the moment showing the largest LV cavity, and end-systole as the moment showing the smallest LV cavity. As no arrhythmia was detected, heart rate (HR) was assessed by calculating the time interval between two consecutive maximal systolic excursions of the LV myocardial walls. Vertical anechoic artifacts (**Fig. 3**) were frequently observed in all elephants and were likely related to their integumentary characteristics, including thick and irregular skin and coarse hair shafts, a feature not encountered in small carnivores. However, these artifacts did not interfere with the acquisition of measurements. Spontaneous echo contrast was also commonly observed, particularly in the left parasternal long-axis view (**Fig. 3**). This finding was likely related to the combination of low HR and the large size of the LV cavity, both of which promote low-flow conditions, further accentuated by the relatively high gain settings required for deep thoracic imaging.

**Fig 3:**
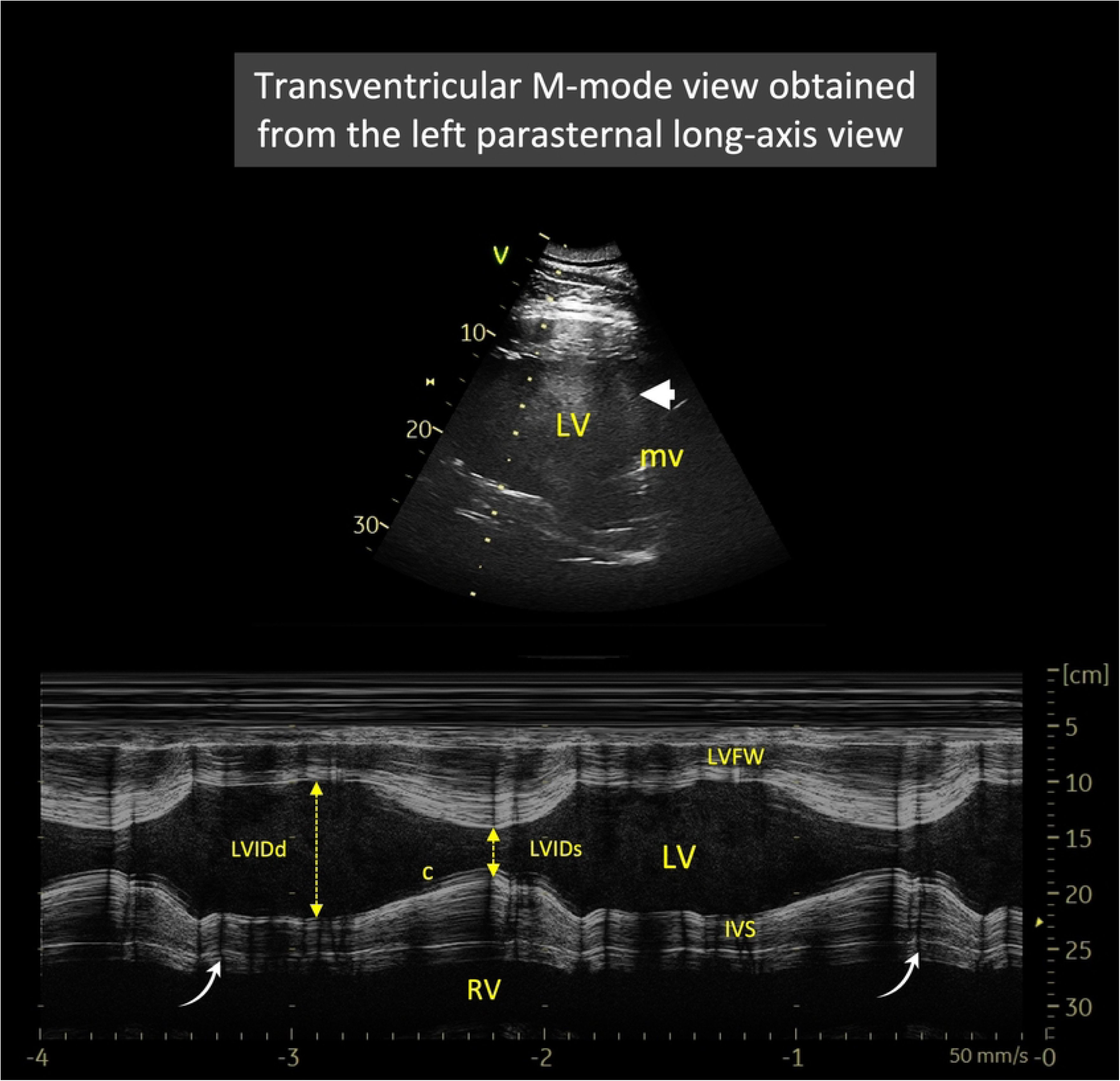
Representative transventricular M-mode echocardiogram obtained from the left parasternal long-axis view in one Asian elephant (*Elephas maximus*) from the study. This view shows two complete cardiac cycles (heart rate = 32 bpm), with the left ventricular free wall (LVFW) proximally and the interventricular septum (IVS) distally, delineating the left ventricular (LV) cavity. The M-mode cursor is positioned perpendicular to the LV myocardial walls, immediately below the insertion of the papillary muscles to exclude them from the measurements. Note the presence of chordae tendineae (c), which were not included in the diameter measurements. A spontaneous echo contrast is typically observed (arrowhead). In addition, vertical anechoic artifacts (curved arrows), corresponding to integumentary shadowing artifacts, are also seen (see text for explanation). *LVIDd and LVIDs: left ventricular internal diameter in end-diastole and end-systole, respectively. RV: right ventricle. Image by Valérie Chetboul*.

### Left parasternal short-axis view and corresponding two-dimensional measurements (Figs. 2 and 4)

The left parasternal short-axis view was obtained by keeping the transducer in the same intercostal space as for the long-axis view and by rotating it by approximately 70-90°, depending on the individual, resulting in a horizontal or nearly horizontal orientation (**Fig. 2**). This view displayed the LV cavity in cross-section with its two papillary muscles (**Fig. 4**), an orientation that differed from the conventional short-axis transventricular view typically obtained from the right parasternal approach in domestic carnivores [5]. Using this view, the internal area of the LV was traced at end-diastole and end-systole (**Fig. 4**), with the papillary muscles excluded from the tracing. The LV FAC% was calculated as the percentage change in LV cross-sectional area between diastole and systole [14]. As no concomitant ECG tracing could be performed, end-diastole and end-systole were defined based on mechanical events; specifically, end-diastole was identified as the frame showing the largest LV cavity, and end-systole as the frame showing the smallest LV cavity.

**Fig 4.**
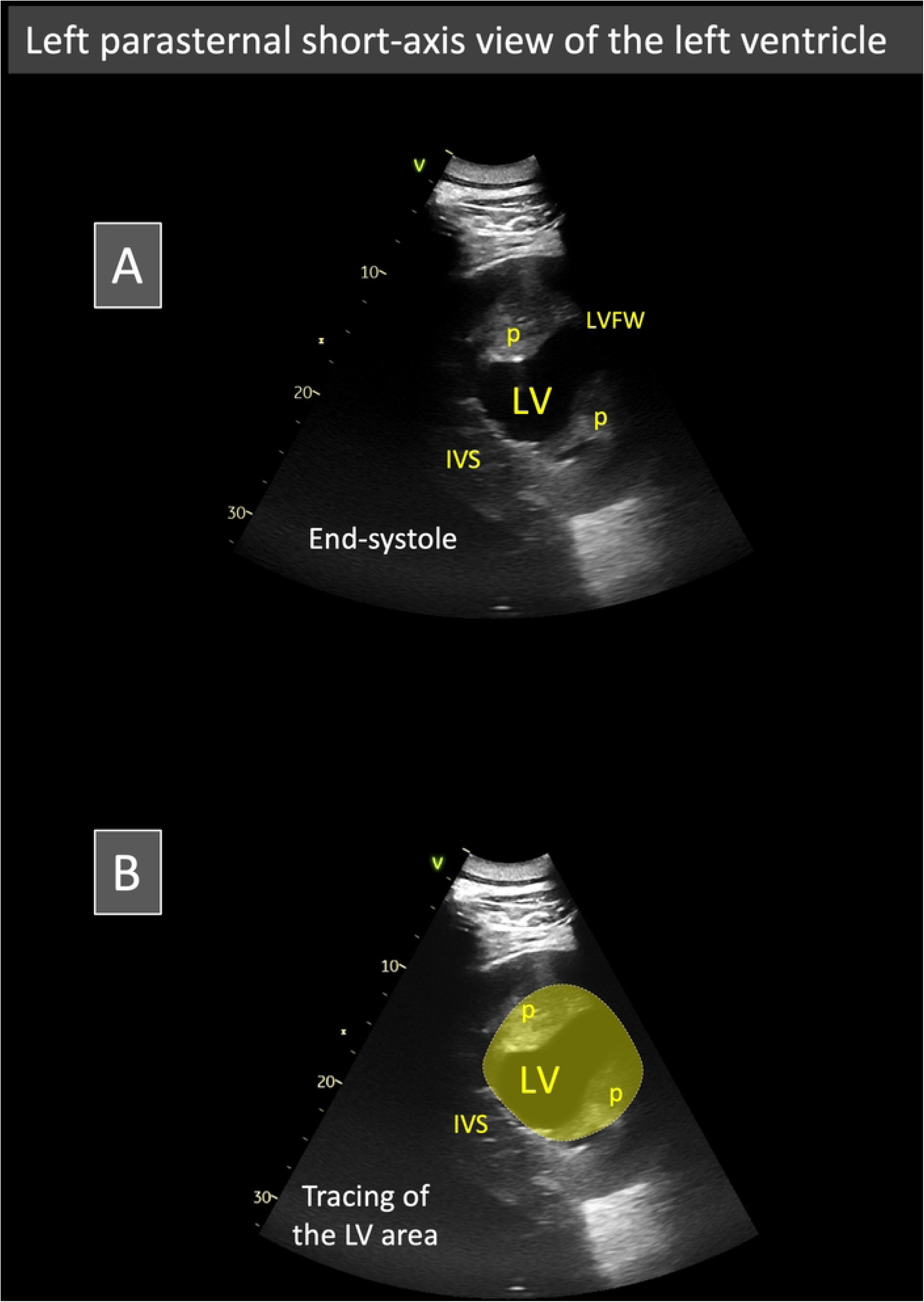
(A and B): Representative two-dimensional left parasternal short-axis view of the left ventricle in one Asian elephant (*Elephas maximus*) from the study at end-systole, showing the left ventricular (LV) cavity in cross-section and the two papillary muscles (p). An example of LV cavity tracing, excluding the papillary muscles, is illustrated in panel B. A similar tracing was performed on the frame obtained at end-diastole, and the LV fractional area change (FAC%) was calculated as [(LV area in diastole – LV area in systole) / LV area in diastole] × 100. *IVS: interventricular septum. LVFW: left ventricular free wall. Image by Valérie Chetboul*.

### Assessment of intra-observer within-day and between-day variability of TTE variables

A total of 72 echocardiographic examinations were performed on 4 different days over a 3-month period in the three elephants EL1, EL2, and EL3, using the above-described echocardiographic procedure. Each elephant was examined 6 times per day. As mentioned above, each TTE examination was performed by the same trained observers: one (Observer 1, AB) holding the transducer and performing the echocardiographic examination, and the other (Observer 2, VC, Dipl. ECVIM-CA Cardiology) adjusting the settings in real time for optimal image acquisition. Each examination included one 2D short-axis view of the LV and one 2D-guided transventricular M-mode view, with offline assessment of eight TTE variables (LVIDd, LVIDs, LVFWd, LVFWs, IVSd, IVSs, SF%, and LV FAC%). A general linear model was used to determine the within-day and between-day coefficients of variation (CV).

### Statistical analysis

All analyses were conducted using RStudio (Version 2023.09.1+494), with the packages lme4, broom.mixed, dplyr, tidyr, writexl, and ggplot2. For each echocardiographic parameter, descriptive statistics (mean, standard deviation (SD), minimum, and maximum) were calculated for the entire dataset as well as separately for each individual elephant. To assess measurement variability, linear mixed-effects models were fitted for each parameter with animal, day, and the animal × day interaction specified as random effects. This allowed partitioning of the total variance into three components: within-day variability (repeatability), between-day variability (reproducibility), and interindividual variability. For each source of variability, the standard deviation was extracted from the model, and the corresponding CV was calculated by dividing the SD by the overall mean of the parameter. This approach provided a normalized estimate of the magnitude of each variance component.

Additionally, the influence of HR, body weight, and age on echocardiographic parameters was investigated using multivariable linear mixed-effects models. For each parameter, HR, body weight, and age were included as fixed effects, while animal identity was modeled as a random intercept to account for repeated measures within individuals. Model estimates, 95% confidence intervals (CI), and P-values were extracted for each predictor. A P-value < 0.05 was considered statistically significant.

## Results

All 72 TTE examinations were successfully performed in the three elephants (EL1, EL2, EL3), and all eight variables, including six M-mode measurements (LVIDd, LVIDs, LVFWd, LVFWs, IVSd, and IVSs) and two derived indices (SF% from M-mode and LV FAC% from 2D mode), were assessable offline, yielding a total of 576 data points for the study. The mean heart rate ± SD during TTE examinations was 40 ± 4 beats/min (range: 32-47).

**Table 1** presents the values of the 576 repeated TTE measurements for the entire cohort and for each elephant individually, and the corresponding within-day and between-day SD and CV values are shown in **Table 2**.

**Table 1.**
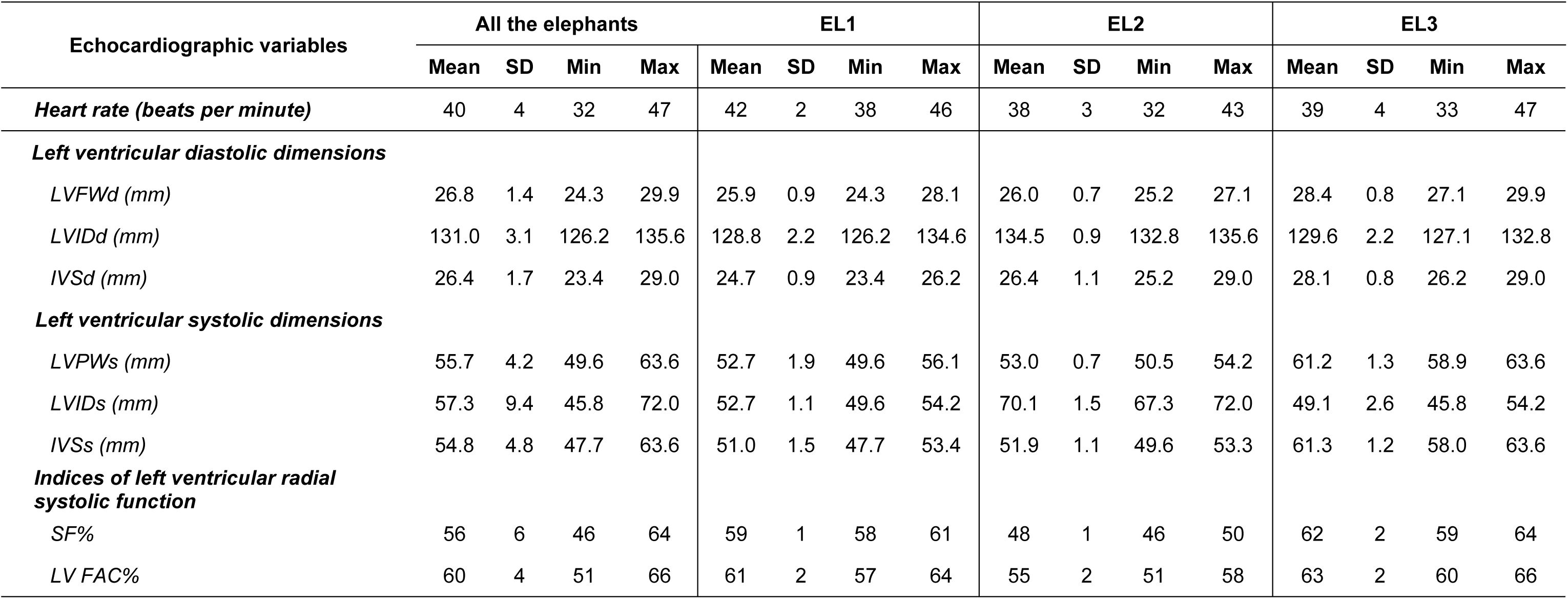
Mean ± standard deviation (SD), minimum (Min), and maximum (Max) values of eight echocardiographic variables obtained from 72 transthoracic examinations performed in three Asian elephants (*Elephas maximus*, EL1–EL3). Results are presented for all elephants combined and individually. The echocardiographic variables included seven repeated M-mode parameters, i.e., interventricular septal thicknesses at end-diastole and end-systole (IVSd, IVSs), left ventricular internal diameters at end-diastole and end-systole (LVIDd, LVIDs) with calculation of the shortening fraction (SF%), and left ventricular free wall thicknesses at end-diastole and end-systole (LVFWd, LVFWs), as well as one two-dimensional parameter, the left ventricular fractional area change (LV FAC%). Heart rate (HR) was also calculated.

**Table 2:** Within-day, between-day, and interindividual variability, expressed as standard deviations (SD) and coefficients of variation (CV), for eight echocardiographic variables obtained from 72 transthoracic examinations in three Asian elephants (*Elephas maximus*). These variables included seven repeated M-mode parameters, i.e., interventricular septal thicknesses at end-diastole and end-systole (IVSd, IVSs), left ventricular free wall thicknesses at end-diastole and end-systole (LVFWd, LVFWs), and left ventricular internal diameters at end-diastole and end-systole (LVIDd, LVIDs) with calculation of the shortening fraction (SF%), as well as one two-dimensional parameter, the left ventricular fractional area change (LV FAC%).

| Echocardiographic variables | Within-day variability |  | Between-day variability |  | Interindividual variability |  |
| --- | --- | --- | --- | --- | --- | --- |
|  | SD | CV (%) | SD | CV (%) | SD | CV (%) |
| ***Left ventricular diastolic dimensions*** |  |  |  |  |  |  |
| *LVFWd (mm)* | 0.2 | 0.7 | 0.8 | 3.0 | 1.4 | 5.2 |
| *LVIDd (mm)* | 0.4 | 0.3 | 1.8 | 1.4 | 3.0 | 2.3 |
| *IVSd (mm)* | 0.3 | 1.2 | 0.9 | 3.4 | 1.6 | 6.2 |
| ***Left ventricular systolic dimensions*** |  |  |  |  |  |  |
| *LVFWs (mm)* | 0.5 | 0.8 | 1.3 | 2.4 | 4.8 | 8.7 |
| *LVIDs (mm)* | 0.9 | 1.6 | 1.7 | 2.9 | 11.2 | 19.6 |
| *IVSs (mm)* | 0.0 | 0.0 | 1.3 | 2.4 | 5.7 | 10.3 |
| ***Indices of left ventricular radial systolic function*** |  |  |  |  |  |  |
| *SF%* | 0.5 | 1.0 | 1.2 | 2.1 | 7.5 | 13.4 |
| *LV FAC%* | 0.0 | 0.0 | 1.9 | 3.1 | 4.3 | 7.2 |

All within-day and between-day CV values (n=16) were < 10% (0 to 3.4%), the lowest being respectively observed for IVSs and LV FAC (0%), and LVIDd (1.4%). Interindividual variability was higher, ranging from 2.3% to 19.6%.

The effect of HR, body weight, and age on echocardiographic parameters was assessed using a multivariable linear mixed-effects model. Age was negatively associated with LVFWd (Estimate = –2.38 [95% CI: –4.64 to –0.12], P = 0.04), LVFWs (–9.46 [–18.37 to –0.55], P = 0.04), and LVIDd (–15.54 [–27.58 to –3.51], P = 0.01), and positively associated with LV FAC% (0.40 [0.40 to 0.50], P < 0.0001). Body weight was positively associated with LVIDd (0.80 [0.23 to 1.37], P = 0.01) and negatively with LV FAC% (–0.02 [–0.02 to –0.01], P < 0.0001). Lastly, HR was not significantly associated with any parameter in the multivariable models. Full results are presented in **Table 3**.

**Table 3.** Associations of heart rate (HR), body weight, and age with echocardiographic variables in Asian elephants (Elephas maximus) included in the study. HR are expressed in beats per minute, body weight in kilograms, and age in years. Echocardiographic dimensions are in millimeters (mm), and indices in percent (%). Regression estimates are expressed per unit increase in the corresponding predictor. Multivariable linear mixed-effects models were fitted for each echocardiographic parameter with HR, body weight, and age as fixed effects and individual animal as a random effect. The table reports the estimated effect size (Estimate), the 95% confidence interval (CI), and the corresponding P-value for each predictor. Significant associations (P < 0.05) are shown in bold. *IVSd and IVSs: interventricular septal thickness at end-diastole and end-systole, respectively. LVFWd and LVFWs: left ventricular free wall thicknesses at end-diastole and end-systole, respectively. LVIDd and LVIDs: left ventricular internal diameter at end-diastole and end-systole, respectively. SF%: shortening fraction. LV FAC%: left ventricular fractional area change*.

| Parameter | Predictor | Estimate | Lower 95% CI | Upper 95% CI | P-value |
| --- | --- | --- | --- | --- | --- |
| **LVFWd** | HR | 1.67 | -3.78 | 7.12 | 0.54 |
|  | Body weight | 0.06 | -0.04 | 0.17 | 0.24 |
|  | **Age** | **-2.38** | **-4.64** | **-0.12** | **0.04** |
| **LVIDd** | HR | 4.67 | -24.36 | 33.71 | 0.75 |
|  | **Body weight** | **0.80** | **0.23** | **1.37** | **0.01** |
|  | **Age** | **-15.54** | **-27.58** | **-3.51** | **0.01** |
| **IVSd** | HR | -3.12 | -8.84 | 2.60 | 0.28 |
|  | Body weight | 0.02 | -0.09 | 0.14 | 0.67 |
|  | Age | -1.58 | -3.95 | 0.79 | 0.19 |
| **LVFWs** | HR | 0.25 | -0.82 | 1.32 | 0.64 |
|  | Body weight | 0.46 | -0.45 | 1.37 | 0.19 |
|  | **Age** | **-9.46** | **-18.37** | **-0.55** | **0.04** |
| **LVIDs** | HR | -2.62 | -15.02 | 9.79 | 0.68 |
|  | Body weight | 0.03 | -0.21 | 0.27 | 0.81 |
|  | Age | -1.39 | -6.53 | 3.75 | 0.59 |
| **IVSs** | HR | 4.87 | -1.70 | 11.44 | 0.14 |
|  | Body weight | 0.05 | -0.08 | 0.18 | 0.46 |
|  | Age | 0.77 | -1.95 | 3.49 | 0.57 |
| **SF%** | HR | -0.07 | -0.17 | 0.02 | 0.14 |
|  | Body weight | 0.02 | -0.07 | 0.10 | 0.57 |
|  | Age | -0.34 | -1.16 | 0.48 | 0.40 |
| **LV FAC%** | HR | -0.08 | -0.23 | 0.06 | 0.25 |
|  | **Body weight** | **-0.02** | **-0.02** | **-0.01** | **<0.0001** |
|  | **Age** | **0.40** | **0.40** | **0.50** | **<0.0001** |

## Discussion

Although echocardiography is considered the gold standard non-invasive imaging modality for the assessment of cardiac structure and function in humans and many animal species, its application in elephants has been extremely limited. To date, knowledge of elephant cardiology has mainly derived from necropsy and histopathological examinations [1,2], with only a few ante mortem investigations focusing on heart rate monitoring using ECG or wearable sensors [15–17], on specific ECG methodologies [8,18], or on circulating cardiac biomarkers such as cardiac troponin I in the context of EEHV infection [7,19]. These studies have provided valuable insights into normal cardiac rhythm and ECG tracings, stress-related HR changes, and myocardial injury, but they remain scarce and most often limited by small sample sizes. To the best of the authors’ knowledge, no study has yet provided standardized, quantitative 2D or M-mode TTE data on cardiac morphology and function in awake elephants, and the antemortem diagnosis of heart diseases in this species thus remains challenging. This pilot study represents a critical step toward establishing a reliable diagnostic tool for the antemortem evaluation and longitudinal monitoring of left-sided cardiac disease in Asian elephants. For the first time, we demonstrate the feasibility of a standardized, repeatable, and reproducible TTE protocol in this species. Performing TTE in awake elephants opens new avenues for veterinary care, including pre-anesthetic evaluation, routine monitoring, early detection of subclinical cardiac disease, longitudinal assessment of disease progression, and evaluation of therapeutic interventions, thereby ultimately supporting the conservation of this threatened species. Additionally, integrating TTE monitoring into the management of Asian elephants at risk for, or affected by, EEHV could yield crucial insights into cardiac involvement and complement cardiac troponin I measurement in the overall clinical evaluation of this viral disease [7,19].

Until now, the limited development of TTE in elephants has primarily reflected the substantial technical challenges inherent to this species. Their massive body size complicates the rapid identification of acoustic windows, while the depth and large size of the heart, combined with its relatively cranial position beneath the forelimbs, restrict acoustic accessibility and permit only partial visualization of the organ with currently available equipment [1]. Moreover, the unusually thick and relatively dry skin, together with abundant subcutaneous tissue, further attenuates the ultrasound beam. In addition to these anatomical and technical limitations, the behavioral challenges and potential risks for operators when handling awake elephants have further constrained the use of echocardiography in this species [1]. Collectively, these factors have long hindered the advancement of TTE in elephants. In the present study, several of these obstacles were successfully addressed through specific methodological refinements. The exclusive use of a left parasternal approach proved critical, as right parasternal imaging consistently failed to provide interpretable views owing to the thoracic conformation of the species. Careful preparation of the acoustic window (i.e., soaking the thick skin with lukewarm water, allowing sufficient time for softening, and then applying abundant ultrasound coupling gel) was also essential to ensure adequate image quality. The use of a convex 4C-RS probe further expanded the ultrasonic field compared with a phased-array transducer, while its relatively broad footprint fitted well within the wide intercostal spaces of elephants. In addition, optimized gain settings were required to compensate for both the considerable thoracic depth of the heart and the attenuation caused by thick skin and subcutaneous tissue. Extending the left forelimb over a training ball also improved access to the cranial portion of the left thoracic wall, thereby facilitating the acquisition of both long-axis and short-axis views of the LV. Collectively, these methodological steps, although simple, represent key innovations that enabled us to overcome the technical challenges inherent to TTE in elephants.

Importantly, in this study the TTE protocol was developed through cooperative training and positive reinforcement, allowing elephants to participate voluntarily and remain calm throughout the procedure. This strategy avoided the need for sedation or physical restraint, both of which carry substantial risks in this species and are incompatible with repeated follow-up examinations. By relying on voluntary participation, the protocol is consistent with international animal welfare and safety guidelines [12,13] and illustrates how advanced diagnostic procedures can be incorporated into routine management without compromising the animals’ well-being. The absence of stress during the procedure was further supported by HR values recorded during examinations (mean = 40; range 32 - 47 beats/min), which remained within or near the upper end of values reported for non-stressed Asian elephants [17], even though the elephants examined here (7, 9, and 13 years old) were sub-adults, an age class in which slightly higher resting HR have been reported compared to fully mature adults. Nevertheless, HR variability was not assessed in the present study, and its evaluation would be of particular interest, as it is considered a more sensitive indicator of emotional state than HR alone [15].

The within-day variability of the TTE measurements was minimal, with very low SD and CV across all parameters (CV range: 0.0 to 1.6%), as was the between-day variability (CV range: 1.4 to 3.4%), together indicating high repeatability and reproducibility of both 2D and M-mode measurements (well below the 10% threshold) and demonstrating strong measurement consistency across sessions. By contrast, interindividual variability was more pronounced, with CVs ranging from 2.3% (LVIDd) to 19.6% (LVIDs), reflecting a substantial degree of physiological variation between elephants. Although the animals involved in the study were classified as sub-adults [17], they were at different stages of growth, as indicated by their distinct body weights (1,800 to 3,600 kg) and shoulder heights (2.02 to 2.51 m). This morphological heterogeneity likely accounts for much of the observed interindividual variation.

In the present study, several associations were observed between body weight or age, and TTE parameters using a multivariable linear mixed-effects model. As example, body weight was positively associated with LVIDd, which is consistent with the well-established scaling effect of body size on cardiac dimensions in other species [20,21]. Conversely, body weight was negatively associated with LV FAC%, suggesting that heavier elephants may have relatively lower systolic indices. Nevertheless, these findings must be interpreted with caution given the small sample size, but they highlight the importance of considering morphometric variables when interpreting echocardiographic data in elephants.

This preliminary work on TTE in Asian elephants presents several limitations. Quantitative measurements were restricted to the LV, as this chamber was the most consistently detectable and analyzable from the left parasternal approach. The left atrium, aorta, and right heart chambers could only be partially visualized and were therefore not assessed. Although a large number of repeated examinations (72) and measurements (576) were performed, the small number of animals included represents another limitation of the present study. Furthermore, due to the prominent papillary muscles in the elephant LV, the positioning of the M-mode cursor in the left parasternal long-axis view differed from what is typically performed in small animals [5]: it was placed more apically, at the level of papillary muscle insertion, to avoid their inclusion in the measurements (rather than just below the tips of the mitral leaflets at their maximal diastolic opening). Another limitation is the lack of simultaneous ECG recording, which precluded a precise temporal definition of end-diastole and end-systole. Lastly, as all elephants included in this study were cooperative and accustomed to handling, the results presented here may not be directly applicable to non-trained or wild individuals. Moreover, as previously demonstrated by our group in dogs and cats [22,23], TTE is a highly observer-dependent examination. Therefore, the present results are valid only for the two observers involved (Observer 1 and Observer 2). The authors thus encourage veterinarians to determine their own intra- and inter-observer variability before undertaking further echocardiographic studies in elephants.

## Conclusions

In conclusion, this study demonstrates that TTE can be successfully performed in awake captive Asian elephants following a relatively short period of cooperative training. The protocol established here enabled the assessment of eight 2D and M-mode variables with excellent repeatability and reproducibility, while requiring only minimal restraint. Future studies on larger populations of Asian elephants are now warranted to establish reference intervals for these TTE parameters, which will be essential for detecting pathological changes in chamber size and wall thickness, as well as for the diagnosis of LV dysfunction. This work lays the foundation for establishing TTE as a non-invasive and non-stressful diagnostic tool in elephants, representing a critical step toward integrating it into routine preventive health care and improving both clinical care and the long-term conservation of this threatened species.

## Acknowledgements

The authors sincerely acknowledge the “Fondation Un Cœur” (a foundation under the aegis of the “Fondation de France”) for sponsoring the internship of Dr. Morgan Bureau at the Ménagerie, the Zoo of the Jardin des Plantes (Muséum National d’Histoire Naturelle, Paris, France). The authors also warmly thank *La Tanière – Zoo Refuge* (28630 Nogent-le-Phaye, France) for their availability and kind welcome, and in particular Sébastien Muller (Zoological Director / Licensed Curator), Joss Graffin (Co-Manager, Elephant Section), Valentin Larret (Co-Manager, Elephant Section), Amélie Boulay (Keeper), and Maxime Fournier (Keeper).

